# Amplification lags nonlinearity in the recovery from reduced endocochlear potential

**DOI:** 10.1101/2020.05.11.089789

**Authors:** C. Elliott Strimbu, Yi Wang, Elizabeth S. Olson

## Abstract

The mammalian hearing organ, the cochlea, contains an active amplifier to boost the vibrational response to low level sounds. Hallmarks of this active process are sharp location-dependent frequency tuning and compressive nonlinearity over a wide stimulus range. The amplifier relies on outer hair cell (OHC) generated forces driven in part by the endocochlear potential (EP), the ~ +80 mV potential maintained in scala media, generated by the stria vascularis. We transiently eliminated the EP in vivo by an intravenous injection of furosemide and measured the vibrations of different layers in the cochlea’s organ of Corti using optical coherence tomography. Distortion product otoacoustic emissions (DPOAE) were monitored at the same times. Following the injection, the vibrations of the basilar membrane lost the best frequency (BF) peak and showed broad tuning similar to a passive cochlea. The intra-organ of Corti vibrations measured in the region of the OHCs lost their BF peak and showed low-pass responses, but retained nonlinearity, indicating that OHC electromotility was still operational. Thus, while electromotility is presumably necessary for amplification, its presence is not sufficient for amplification. The BF peak recovered nearly fully within 2 hours, along with a non-monotonic DPOAE recovery that suggests that physical shifts in operating condition are a final step in the recovery process.

**SIGNIFICANCE:** The endocochlear potential, the +80 mV potential difference across the fluid filled compartments of the cochlea, is essential for normal mechanoelectrical transduction, which leads to receptor potentials in the sensory hair cells when they vibrate in response to sound. Intracochlear vibrations are boosted tremendously by an active nonlinear feedback process that endows the cochlea with its healthy sensitivity and frequency resolution. When the endocochlear potential was reduced by an injection of furosemide, the basilar membrane vibrations resembled those of a passive cochlea, with broad tuning and linear scaling. The vibrations in the region of the outer hair cells also lost the tuned peak, but retained nonlinearity at frequencies below the peak, and these sub-BF responses recovered fairly rapidly. Vibration responses at the peak recovered nearly fully over 2 hours. The staged vibration recovery and a similarly staged DPOAE recovery suggests that physical shifts in operating condition are a final step in the process of cochlear recovery.

## INTRODUCTION

In the mammalian cochlea, the sound signal in the form of traveling pressure+motion waves, is converted to electrical signals by the sensory hair cells that lie within the the organ of Corti (Fig.1). Arrays of tightly-packed stereocilia (hair bundles) of graded height protrude from the apical surface of the hair cells. During hearing, relative motion between the tectorial membrane, an acellular structure that overlies and couples the hair cells, and the reticular lamina located at the apical surface of the hair cell bodies, results in shearing of the hair bundles. When the stereocilia deflect towards/away from their tallest row, mechanically gated ion channels on the shorter rows are opened/closed and the flow of cations, mostly K^+^, also Ca^2+^, is increased/reduced and depolarizes/hyperpolarizes the hair cells. Complex sounds are resolved into individual frequency components which peak at different locations along the length of the cochlea with high frequencies encoded at the base and low frequencies encoded at the apex.

The mechanical vibrations of the organ of Corti Complex (OCC = the organ of Corti and tectorial and basilar membranes, TM and BM) are boosted by the cochlear amplifier, an active process or set of coupled processes operating under feedback. A key component of amplification is generated by the outer hair cells (OHCs): in response to changes in the membrane potential, the OHC soma changes length in a process driven by the motor protein prestin (1). The cochlear amplifier increases the response to low level sounds, resulting in a compressive nonlinearity that boosts the dynamic range of hearing across some six orders of stimulus pressure magnitude (2–4). In healthy cochleae, the vibrations at low sound pressure levels are sharply peaked at each longitudinal location’s best frequency (BF) (Fig. 1C and D). Thus the cochlear amplifier increases both sensitivity and frequency resolution. In cochleae for which the active process is not functional, for example: *post mortem*, following mechanical damage, or pharmacological inhibition, the responses become linear and exhibit broad tuning. Broad tuning is also typically observed at stimulus levels greater than about 80 dB SPL. OHC electromotility is most likely responsible for amplifying the OCC vibrations, while the nonlinearity is due to saturation in mechano-electric transduction (MET) (Fig. 1E and F) (5, 6). The nonlinearity of the cochlear amplifier produces distortion product otoacoustic emissions (DPOAEs). When two or more tones are presented simultaneously, additional frequencies are generated in the cochlea and reverse propagate to the middle ear. The resulting motion of the eardrum then produces faint sounds at these distortion frequencies that can be measured in the ear canal. DPOAEs are used as a non-invasive gauge of cochlear condition, and have also been used to detect operating point shifts in the MET nonlinearity (Fig. 1E) (7, 8).

**Figure 1:**
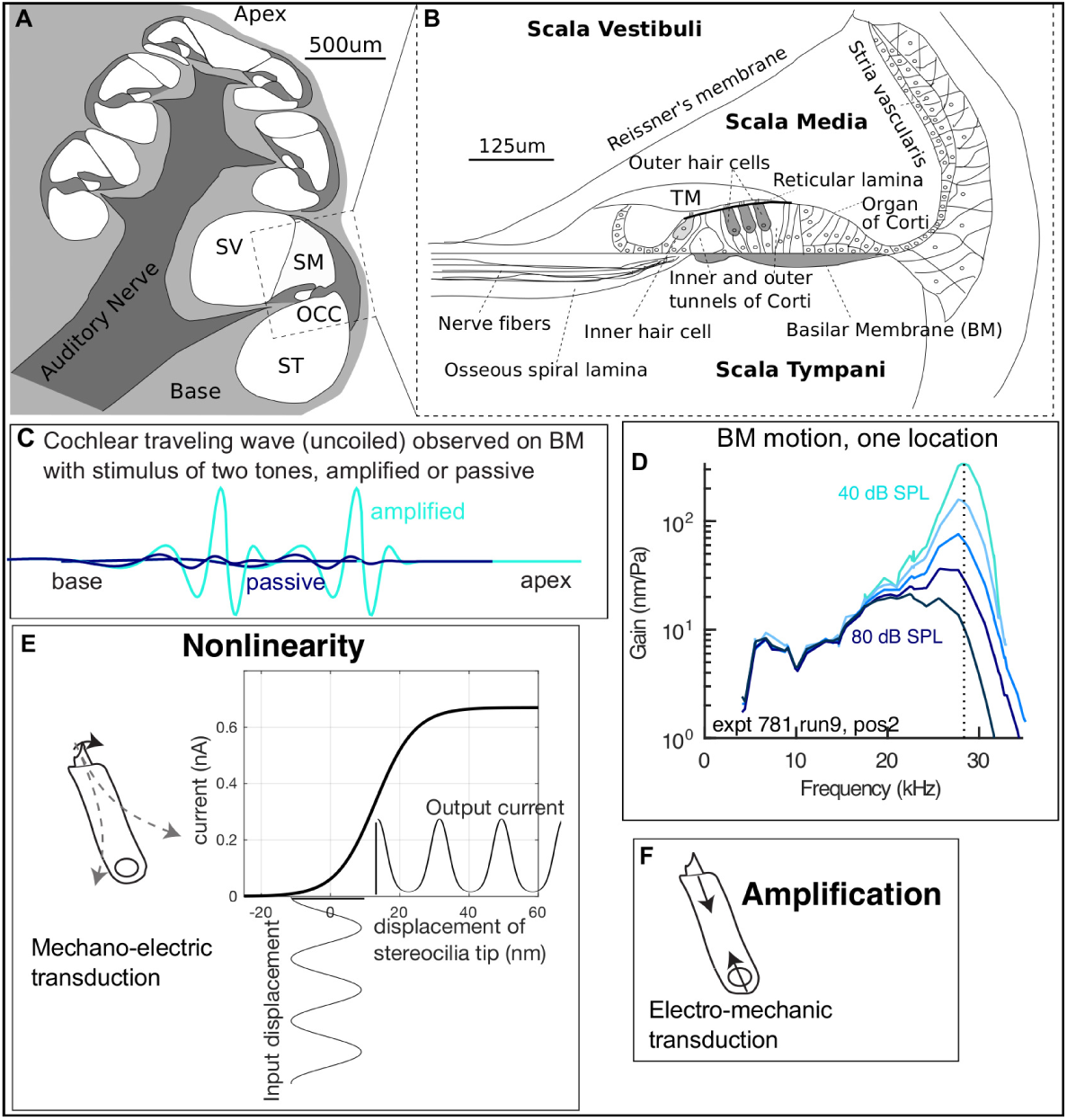
(A) Cross-sectional sketch of the gerbil cochlea, with fluid compartments scala media (SM), scala tympani (ST) and scala vestibuli (SV) labelled. (B) The boxed section in A is expanded, to show the organ of Corti complex (OCC). The OCC is composed of the sensory tissue of the OC (including the inner and outer hair cells (IHC and OHC)) and surrounding acellular structures of the basilar and tectorial membranes (BM and TM). (C) Cartoon of the motion response to two tones on the BM, showing the characteristic amplification in the region where the responses peak, and illustrating that amplification will make otherwise undetectable signals detectable and enhance frequency resolution. (D) BM motion data in a healthy cochlea, showing the hallmarks of amplification: nonlinearity and sharp tuning at low to moderate stimulus levels. The best frequency (BF) is indicated by the vertical dashed line. (E) Nonlinearity is based in the saturation of OHC current when MET channels are pushed to the nearly fully open and fully closed states. OHC current data and Boltzmann function fit redrawn from (6). (F) The mechanical basis of amplification is most likely OHC somatic forces arising from the prestin molecule’s conformation changes following electro-mechanic transduction.

The MET currents are driven in part by the endocochlear potential (EP), the ~ +80 mV electrical potential within the scala media. The EP is generated by cells within the stria vascularis (Fig. 1B). Gradual degradation of the stria vascularis and the corresponding decline in the EP is one cause of presbyacusis, or age-related hearing loss, which affects up to half of the population over 75 years of age in the United States (9–11). Loop diuretics such as furosemide cause a sudden and reversible decrease in the EP and have been used *in vivo* in animal studies to investigate the effects of EP reduction on mechanical (12) and electrical (13–15) responses within the cochlea, and also on DPOAEs (15–17).

In recent studies by our group (15), furosemide was administered intravenously (iv) to gerbil, and EP was monitored continuously along with the extracellular voltage measured close to the BM. We termed this voltage "local cochlear microphonic" (LCM). The LCM shows tuning and traveling wave phase excursion similar to the adjacent BM motion, and is useful for exploring cochlear amplification (18–20). LCM is a measure of OHC current and its saturation can be used as an in-vivo probe of saturation and operating point shifts in MET (Fig. 1 E) (15, 20). In the furosemide + LCM experiments, following a deep reduction in EP and LCM responses, both recovered but with different time scales. Interestingly, LCM could recover fully with EP still substantially sub-normal, contrary to modeling predictions (21). These points are illustrated in Fig. 2, which shows time-variation following iv-furosemide of: (A) EP for five preparations; (B) examples of LCM at the BF for two preparations; (C) MET operating point. Fig. 2A shows that following iv-furosemide, EP recovered and stabilized at a sub-normal voltage by ~40 mins; we assume that a similar time course holds in the current study, which used an identical iv-furosemide protocol. (For technical/practical reasons, EP could not be simultaneously measured in the current experiments.) In the LCM-recovery data of expt. 696 (data with crosses in Fig. 2B) at 40 mins LCM was in an early phase of recovery. There was a boost of recovery at ~70 mins and full recovery at 100 mins, even though EP had stabilized at only 60 mV (expt. 696 in panel A). In the LCM-recovery data of expt. 705 (solid line data in Fig. 2B) at 40 mins LCM had recovered somewhat and nearly plateaued, then had a boost of recovery at 50-80 mins. By analyzing the fundamental and harmonics in expt. 705, MET operating point was determined and its time dependence is shown in Fig. 2C. The study concluded that the delayed LCM recovery coincided with an operating point shift in MET. To summarize the findings of the LCM studies: The recovery of cochlear amplification occurred many mins after the EP had stabilized at a sub-normal level, and that recovery was concurrent with a recentering of MET operating point.

**Figure 2:**
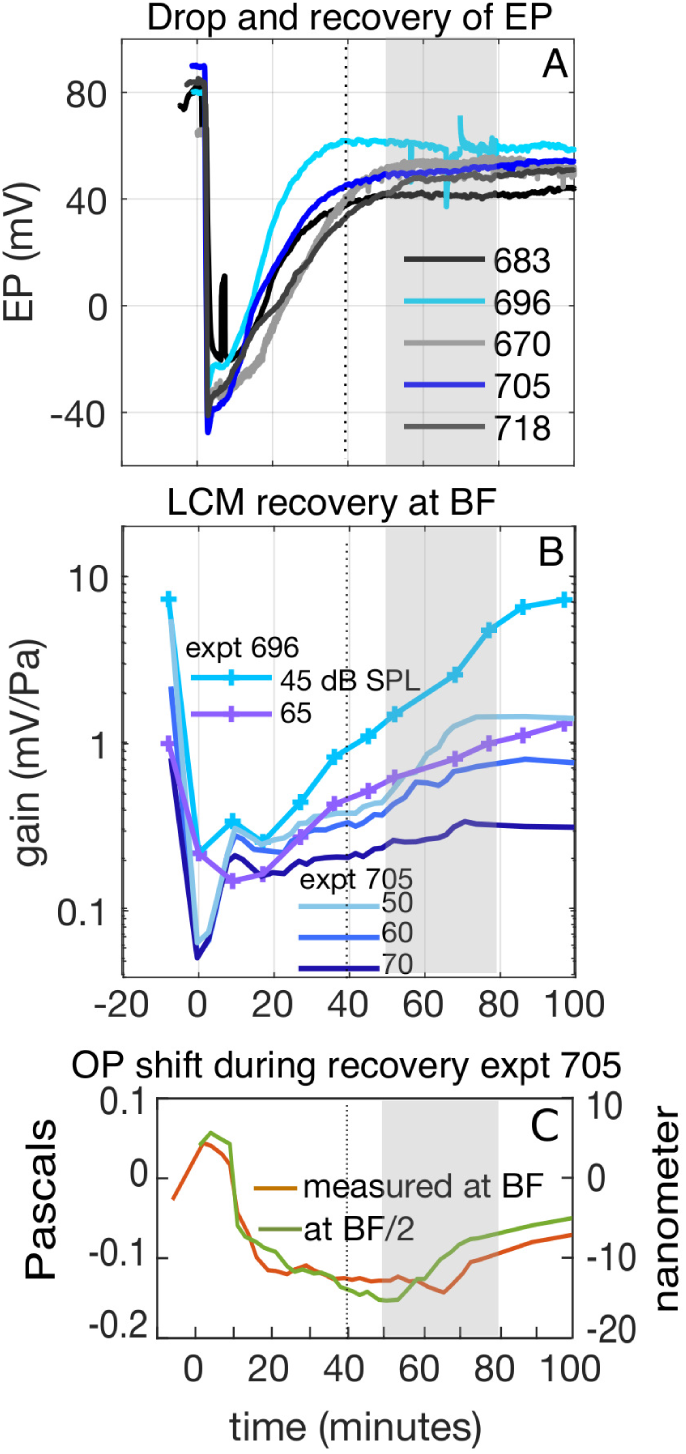
Background results from a recent iv-furosemide study, redrawn from (15). (A) Following iv-furosemide, EP dropped immediately to −20 – −40 mV, began recovering within 10 mins and stabilized at a sub-normal level ~40 mins after furosemide delivery (vertical dotted line). The two blue traces correspond to the experiments in (B). (B) Recovery of LCM responses at BF. LCM recovered more slowly than EP, and underwent a substantial recovery in a 50-80 min time frame where EP had fully stabilized but at a sub-normal level (gray rectangles). In expt. 705 LCM responses were measured at relatively dense time points. (C) MET operating point shifts measured in expt. 705, using second harmonic LCM responses. Left axis corresponds to stimulus pressure in the ear canal, right axis is approximate stereocilia bundle shift. Responses at two stimulus frequencies are shown to demonstrate similarity across frequency. The operating point returns towards zero in the 50-80 min time frame coinciding with the recovery of BF LCM in (B). The size and timing of this operating point variation are useful background information for analyzing the vibration and DPOAE results of the present paper. ((15) used single-tone stimuli; LCM data with multi-tone stimuli (but without iv-furosemide) is in (20).)

Following that background, in the present study we used phase sensitive spectral domain optical coherence tomography (SD-OCT) (22) to measure the sound-induced vibrations within the OCC *in vivo* before and after the EP was perturbed by an iv injection of furosemide. Unlike single-point interferometry which only yields a vibration measurement at one position in the optical path, typically the BM when recording from basal locations in the cochlea (2), OCT allows for simultaneous measurements at multiple axial locations. This enables simultaneous recording from different layers within the organ of Corti, and the ability to contrast the motion at the BM and OHC-regions. To give a brief preview of findings: BM vibrations lost their BF peak and became nearly passive, similar to findings in previous studies (12). The vibrations in the OHC region lost the BF peak and became low-pass, but retained sub-BF compressive nonlinearity. This vibration nonlinearity is taken to be an expression of OHC-electromotility, an expectation that is supported by the observation that the LCM, representing OHC current (and thus voltage), shows substantial sub-BF nonlinearity when elicited with the multi-tone stimuli used in the present study (20). The DPOAEs recovered nonmonotonically, with a time course that reinforced the correlation of MET operating point to the recovery of amplification. Thus, the present findings indicate that (1) the presence of even robust OHC electromotility is not sufficient for effective amplification and (2) effective amplification returns along with a recovery of MET operating point. These findings advance our understanding of the constellation of factors that sustain the cochlear amplifier and produce the remarkable sensitivity and frequency resolution of the cochlea.

## MATERIALS AND METHODS

The experiments were approved by the Columbia University Institutional Animal Care and Use Committee.

### Gerbil Preparation

Young, mature gerbils of either sex were anesthetized with intraperitoneal (IP) injections of ketamine (3–6 mg) and sodium pentobarbital (40 mg/kg). Anesthesia was maintained with supplemental doses of pentobarbital given if the animals displayed a reflex in response to a light toe pinch. Buprenorphone (0.2 mg/kg) was administered IP every six hours. The gerbil’s scalp was removed and the head attached to a two-axis goniometer (Melles-Griot) with dental cement (Durelon, 3M). The left pinna and most of the cartilaginous ear canal were resected and the animals were tracheostomized to facilitate breathing. The tissue and muscle over the left temporal bone were carefully dissected and a narrow opening in the bulla was made by chipping the bone with fine forceps. A bridge of dental cement was used to firmly attach the bulla to the goniometer. Throughout the surgery and experiment, the animal’s temperature was maintained at 38°C with a servo-controlled heating blanket and monitored with a rectal thermometer. During the OCT measurements, additional heating to the animal’s head was supplied by a disposable hand warmer (Hot Hands, HeatMax Inc.) positioned on the goniometer. Experiments were conducted on an optics table in an acoustical isolation booth (Industrial Acoustics Corp.).

### Acoustical System

Acoustic signals were generated using a Tucker Davis Technologies (TDT) system with a sampling rate of 97656.25 S/s. The sound was played by a Radio Shack speaker and delivered closed-field to the ear canal through a plastic tube. Pressures were measured with an ultrasonic microphone (Sokolich) whose probe tube was positioned 1 – 2 mm from the tympanic membrane. Sound pressure levels are reported as dB SPL referenced to the standard value, 0 dB = 20 *µ*Pa. The OCT and TDT systems were synchronized as previously described (22). The clock signal from the TDT zBus was modified using a custom built digital/analog circuit to give a high duty cycle (90% high, 10% low) 5 V square wave which served as the TTL trigger for the OCT’s line camera. Tuning curves were measured in response to zwuis tone complexes (23, 24); 60 frequencies from ~ 4 kHz to ~ 37 kHz were presented simultaneously for 2^20^ = 1048576 shots or 10.7 s at 40 to 80 dB SPL in 10 dB steps. The frequencies were chosen such that each component contained an integer number of points per cycle and the stimulus contained no harmonics or distortion products up to third order (24). Each sinusoidal component in the complex was assigned a random phase so the pressure magnitude of the complex was ~ √60 higher than the magnitude of each component. DPOAEs in response to swept two-tone (*f*_1_ and *f*_2_) stimuli were measured before each set of tuning curves throughout the experiment. In these measurements, *f*_2_ was varied from 1 to 48 kHz, *f*_1_ and *f*_2_ were held at a fixed ratio of *f*_2_ = 1.2 *f*_1_, and the two primary tones were presented at 50 and 70 dB SPL for 1 s and averaged 50 times. The noise level was typically −3–0 dB SPL for these recordings.

### Optical Coherence Tomography and Spectral Domain Phase Microscopy

Cochleae were imaged with a ThorLabs Telesto III OCT equipped with an LSM03 5×, 0.055 NA objective lens. After the initial animal surgery, the gerbils were placed under the OCT and the instrument’s video camera and an operating microscope were used to position the head. Recordings were made in the cochlear base near the 25 kHz location, found by aiming the system apically through the round window. Initially, continuous two-dimensional scanning was done with the ThorImage program and small adjustments made to the position until the OCC was centered in the field of view. The position was adjusted until the two gaps in the OC corresponding to the inner and outer tunnels (Fig. 1 B) were visible in the B- and A-scans. For vibrometry, the OCT was controlled with custom software written in C++ based on the ThorLabs software development kit (SDK). Before and after each set of tuning curves, the OCT acquired 1 mm wide B-scans and the two images were compared to confirm that the sample was stable over the course of the measurement.

For vibrometry, a single A-scan through the BM and OHC region was selected and time-locked A-scan spectra were recorded as the sound played, and then saved to a hard disk as 16 bit raw files for offline analysis. The time series of spectra (termed M-scan) were converted to a series of complex numbers representing reflectivity versus depth (see (22) for analytical details). Pixels of local maxima in the time-averaged A-scan magnitude, corresponding to specific features in the OCT image, were selected for further analysis, for example, a pixel on the round window membrane and several pixels in the BM and OHC region. In spectral domain phase microscopy, the displacement-vs-time of each pixel in the A-scan is proportional to the phase-vs-time of the complex M-scan at that pixel. The noise level in the vibration measurement is determined by the magnitude of the selected A-scan pixel, which depends on the feature and reflectivity. In the best preparations, the noise floor could be as low as. 02 –. 05 nm. Noise rises if the A-scan peak is reduced (Fig. 3 A), which can happen due to micrometer-scale shifts in the preparation. Several baseline measurements were made before the furosemide to establish a solid baseline and the most complete data set was used for presentation. The vibration and interleaved DPOAE measurements were stable during these baseline runs; the DPOAE results in Fig. 10 illustrate that stability.

**Figure 3:**
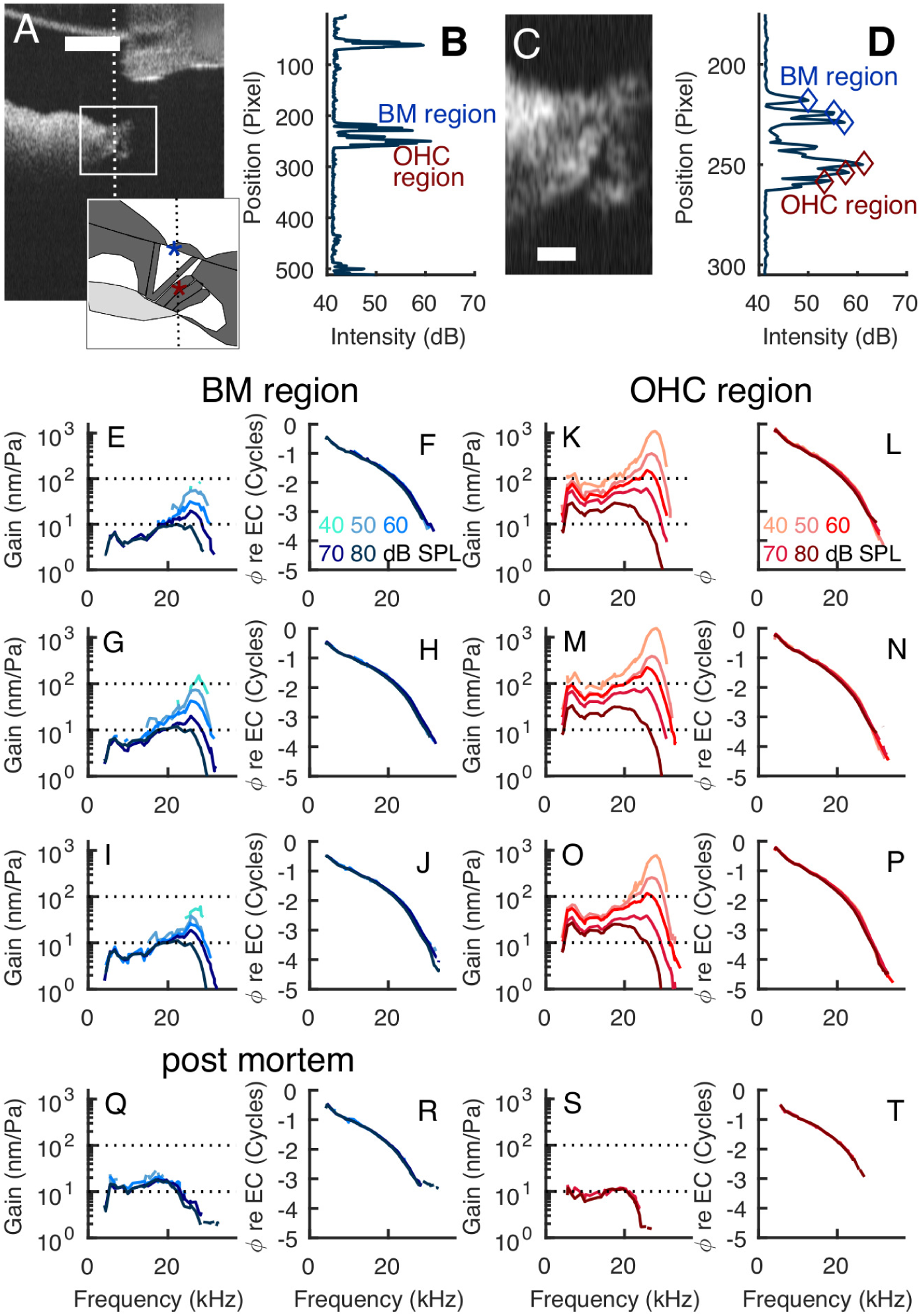
(A) B-scan with sketch showing OHC and BM regions, which were identified using the dark fluid gaps in the B-scan and known anatomy. (Sketch in lower right corner is upside-down and simplified version of Fig.1B. Blue and maroon stars indicate BM and OHC regions.) Scale bar 250 *µ*m. (B) A-scan corresponding to the vertical line in A; this is the line along which vibration measurements were made. The A-scan shows peaks at the round window membrane (top of A-scan) and in the BM and OHC regions. The distance between adjacent pixels is ~3 *µ*m. (C) Expanded view of B-scan. (D) Expanded view of A-scan with locations where vibration was analyzed identified with diamonds. (E-J) Gain and phase re: EC pressure from the three identified points in the BM region. Darker shades indicate higher sound pressure levels, key in panels F and L. K-P Gain and phase: re: EC pressure from the three identified points in the OHC region. (Q-T) *Post-mortem* responses in the two regions. Only the 70 and 80 dB stimuli gave displacements that were significantly above the noise threshold. Expt. 789, 10-2-2019.

Once the vibration time waveforms were acquired for the regions of interest, the amplitudes and phases at the sound stimulus frequencies were extracted by Fourier analysis. For each stimulus frequency, the response was deemed significant if the Fourier coefficient was 3.2 times larger than the standard deviation of the noise level, measured from ten neighboring bins in the spectra. (In the presented plots, when individual data points met that criterion but neighboring points did not, they were usually removed; this removed distracting isolated points that were very close to the defined noise floor.) The resulting tuning curves are presented in terms of the gain, defined as displacement per unit pressure. The phases are referenced to the ear canal pressure. Analysis scripts were custom written in Matlab.

At the start of an experiment, we tested the displacements at one or more locations by playing a short zwuis complex containing 10 frequencies from 10 to 40 kHz at 60, 70, and 80 dB for 1 s. We could perform a complete analysis on these short recordings in approximately five mins and used the resulting coarse tuning curves to gauge the quality of the displacements at the selected location and to estimate the best frequency.

### Experimental Paradigm

After the positioning described above we took a series of baseline measurements each consisting of a set of DPOAE audiograms and vibration tuning curves. After these were completed, the gerbil was given an iv injection of furosemide (100 mg/kg) in the left femoral vein. A set of distortion product measurements was taken immediately after the injection. When the injection was successful, the 2 *f*_1_ – *f*_2_ DPOAEs at 50 dB typically fell to the noise level and the DPOAEs at 70 dB were greatly reduced in amplitude. A set of accompanying zwuis tuning curves was then taken. The measurements were repeated approximately every 10 mins for up to four hours post injection. The DPOAE measurements took approximately three mins each and a complete set of tuning curves took approximately four mins, primarily due to the time required, ~ 40s, to transfer large raw files to a solid state drive. Due to the long time required for each experiment, some drift in the preparation was inevitable. Between recordings, we monitored the position of the cochlea with ThorImage and made minor adjustments to the position as needed. At the conclusion of the experiment, the animals were overdosed with pentobarbital. In some preparations, a set of *post mortem* tuning curves was acquired after the animal expired.

### Electrophysiological Recordings

The EP and microphonic potentials (termed local cochlear microphonic, LCM) were measured as described previously (15), and the results shown here in Fig.2 are redrawn from that study, as they provide important background information. The LCM was measured with an insulated tungsten electrode (tip diameter of ~1 *µ*m) inserted into scala tympani and advanced close to the BM at the base of the cochlea, close to the 18 kHz location. The LCM was measured in response to pure-tone stimuli. The voltages were amplified 500 – 1000x (PARC EG&G) and recorded with the TDT system. The EP was measured with an ~ 10 *µ*m diameter glass micro-electrode with an Ag/AgCl pellet and filled with 0.5 M KCl. The reference electrode was filled with standard saline and was placed on the muscle of the right leg. The DC signal was amplified 10x and recorded every second with a usb DAQ board (DATAQ Instruments Inc.).

## RESULTS

A total of 13 gerbils, 9 male and 4 female, were used in this study. At the start of the measurements, all exhibited large DPOAEs and baseline OCC vibrations were typical of healthy ears. Five of these 13 animals survived at least two hours post iv-injection of furosemide and showed significant, and in three cases nearly complete, recovery of cochlear amplification at the best frequency. Of these five experiments, results from the two that showed the most complete recovery are shown in detail and results from two others is shown in grouped data. The third animal that showed nearly complete recovery suffered from mechanical drift in the apparatus, and thus recovery was not successfully tracked. Four animals showed only partial recovery, and died between 50 and 100 mins after the injection; results from these is shown individually and grouped. In one experiment, the gerbil survived for several hours post-injection and the vibrations showed a decreased amplitude following furosemide and robust recovery in the sub-BF region but an irreversible loss of amplification at frequencies close to BF. The observed incomplete recovery in several preparations is not unexpected given the invasive nature of the surgery and the long time scale of the experiment. Finally, in three experiments, the iv injection was not successful.

The primary measurement of this study was that of vibration within the OCC during recovery from furosemide-induced reduction of EP. Sound stimuli consisted of multi-tone “zwuis” stimuli, in which at each amplitude level, all tones are delivered simultaneously. The tuning observed with multi-tone stimuli is similar to that with pure-tone stimuli, but nonlinearity is more pronounced, likely due to the increased sound volume (20, 24). We concentrated on two regions within the OCC, the BM and the OHC region, as shown in introductory Fig. 3. The two regions can be found based on the surrounding fluid-filled regions of the inner and outer tunnels of Corti, which are dark in the B-scan (see Fig. 1B for anatomical labels). The outer tunnel is distinct and the inner tunnel between the pillars is less distinct in this particular B-scan, but still detectable. The vertical line in the B-scan indicates the location of the A-scan where motion was measured, and the SD-OCT technique makes simultaneous motion measurements at all locations in the A-scan. In Fig. 3E-J motion responses are shown from three points within the BM region; in Fig. 3K-P responses are shown from three points within the OHC region. The spacing between BM and OHC-region locations was typically ~ 60 *µ*m. Both BM and OHC regions are tuned and nonlinear, but the OHC-region responses are up to an order of magnitude larger than BM responses, and compressive nonlinearity in the BM responses begins ~ 1/2 octave below the BF, whereas in the OHC-region nonlinearity extends throughout the entire frequency range. These findings are similar to previous measurements obtained by OCT and interferometry (20, 23, 25–27). The intra-region differences in vibration are small compared to inter-region differences. This is an important point because motion results are most reliable when taken from a local maxima in the A-scan (28) and in the hours-long furosemide study the preparation and image underwent small shifts and therefore the exact same points were not probed during the course of the experiment. *Post-mortem* responses are shown in Fig. 3Q-T; both OHC and BM regions became linear and were either tuned broadly (BM) or nearly low-pass (OHC region).

We used furosemide to reversibly reduce the EP as in Fig. 2A and measured OCC vibrations *in vivo* before and after the introduction of the drug. Recordings were taken immediately after the injection and repeated every ten mins for up to four hours. Interwoven with the vibration measurements, DPOAEs elicited with equal-level primaries of 50 and 70 dB SPL were measured. Data from the two preparations with the fullest recovery are presented in Figs. 4 and 5. These show DPOAE-grams and BM and OHC-region tuning curves measured at select time points.

**Figure 4:**
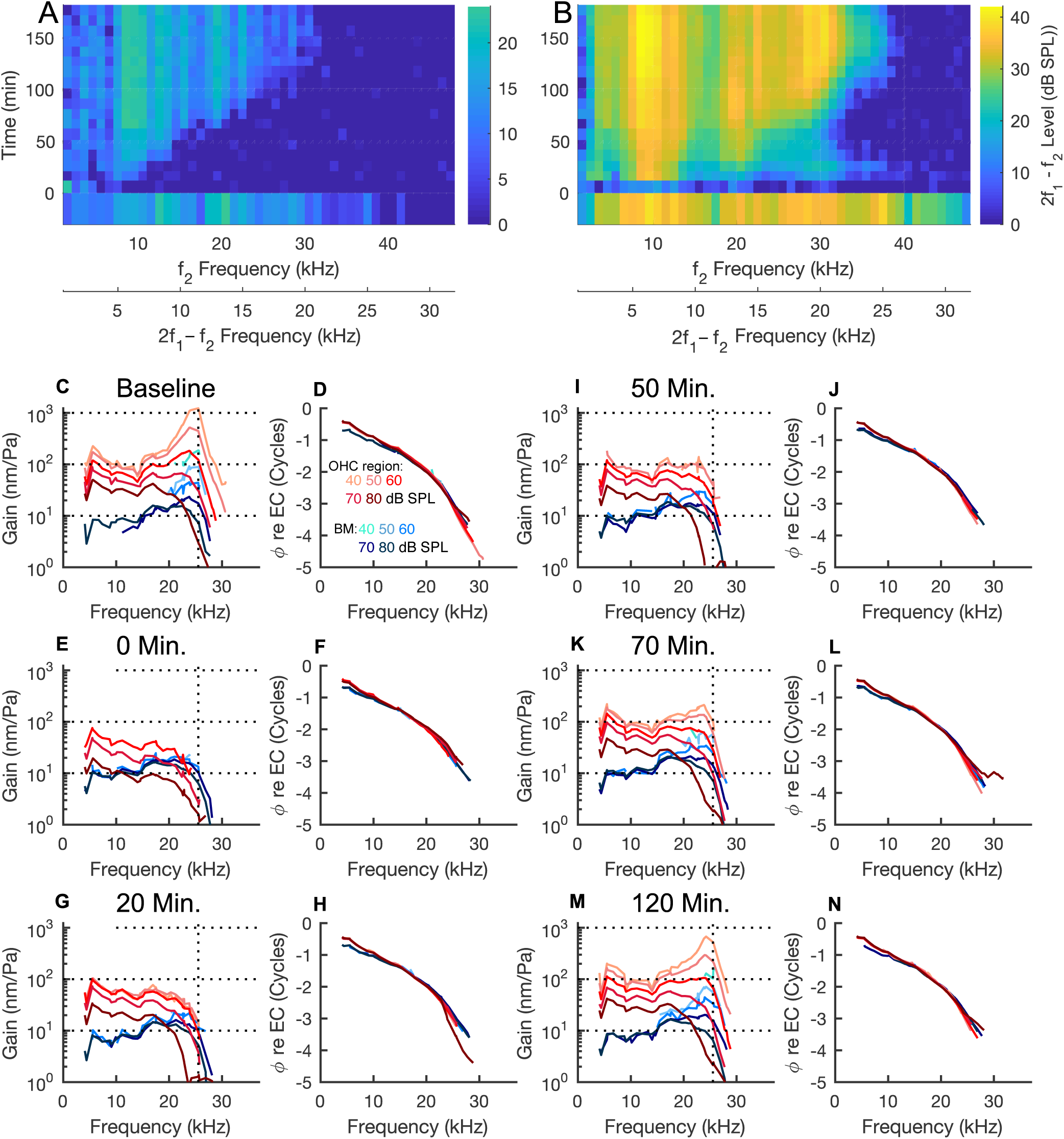
Loss and recovery of amplification following furosemide injection. (A) and (B) show the 2 *f*_1_ – *f*_2_ DPOAE levels evoked by the 50 dB SPL (A) and 70 dB SPL primaries (B). Note the sharp drop immediately following the injection at *t* = 0 mins and the gradual recovery which was maximal at ~ 120 mins. The two x-axes indicate both the 2 *f*_1_ – *f*_2_ and the *f*_2_ frequencies. (C-N) show the displacements and phases measured at the BM and OHC-regions at different time points. BM / OHC-region vibrations are plotted in blue / red with darker shades indicating higher sound pressure levels; key is in panel D. Vibration amplitude is plotted normalized to the stimulus level in the ear canal, and phase is plotted relative to ear canal pressure. (C) and (D) show the baseline responses, recorded before the injection. (E) and (F) were recorded immediately after and the remaining plots show the results at the indicated time points. By 120 mins, the vibrations had recovered to near baseline levels with no further recovery seen thereafter. Expt. 785, 8-28-2019.

**Figure 5:**
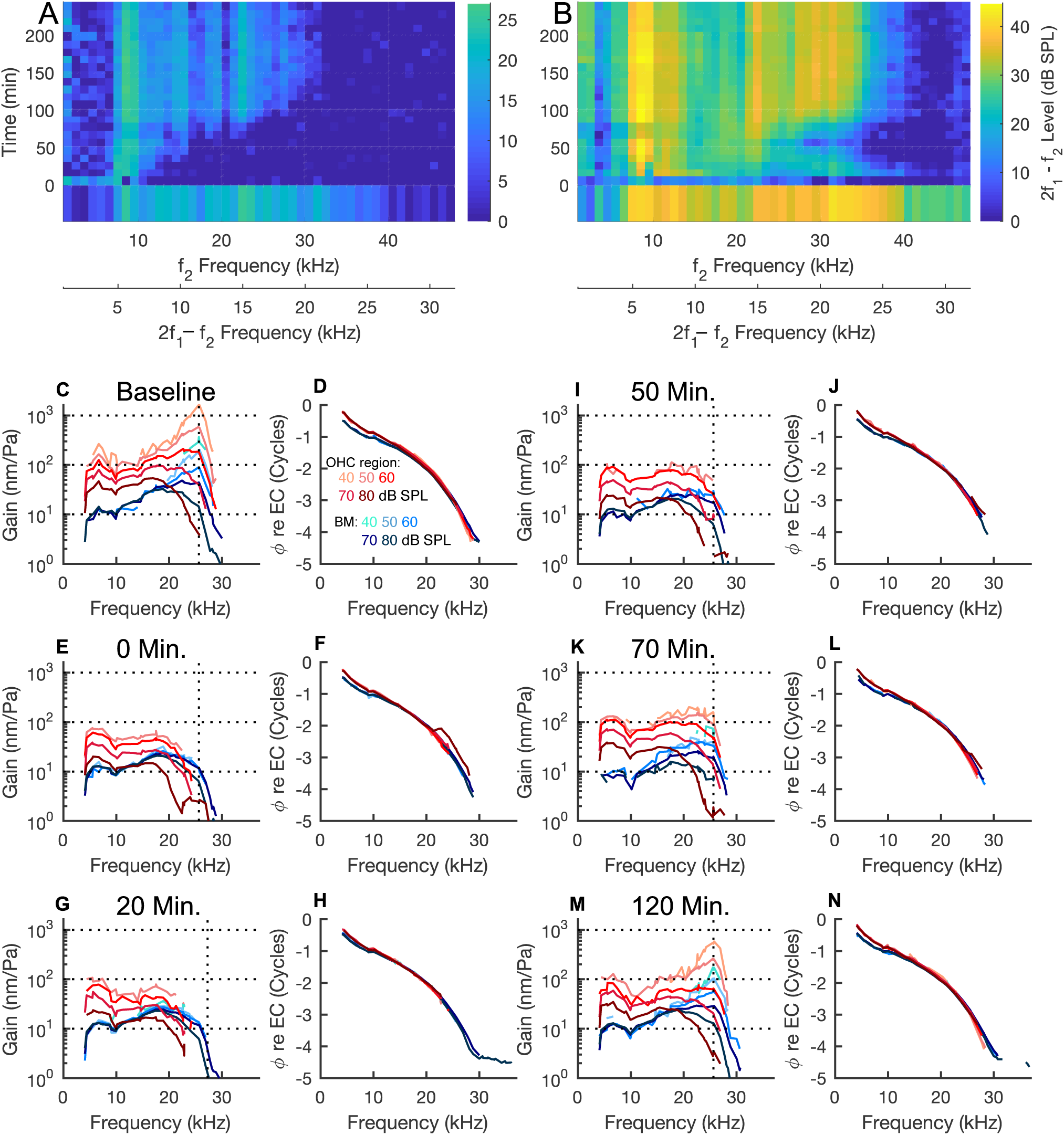
Loss and recovery of amplification in a second cochlea. The layout is as in the previous figure. In this experiment, data were collected for nearly four hours post injection but the recovery of DPOAEs and vibrations reached their fullest extent by about 120 mins post injection and did not recover past that point. Expt. 788, 9-17-19.

Prior to the injection, baseline BM tuning curves and phases were typical of healthy ears (blue curves in C and D of Figs. 4 and 5). The gains showed the characteristic peak at the BF (~26 kHz for both of these preparations) a compressive nonlinearity at nearby frequencies, and linear growth for sub-BF frequencies, starting about 1/2 octave below BF. As in Fig. 3, OHC-region vibrations (red curves) exhibited higher amplitudes, or equivalently higher gains, than the BM vibrations and were compressively nonlinear across the entire frequency range tested. The phases showed traveling wave phase accumulation through more than 4 cycles, with only subtle SPL-dependence. The OHC-region phases led the BM-region phases by ~ .02 cycles at 5 kHz, reducing to ~ .01 cycles at 10 kHz and could lag slightly at frequencies above 20 kHz.

Following the furosemide injection, the BM responses resembled those of a passive cochlea (blue curves in panel E of Figs. 4 and 5). In the BF region there was a loss of amplitude particularly for low and moderate SPL responses, and the gain curves showed broad, nearly linear tuning. The responses at 40 and 50 dB SPL often fell below the noise level. The responses in the linear region below the BF were essentially unaffected by the drug. These observations are similar to previous studies of the effects of furosemide on BM vibrations (12). Following iv-furosemide, the OHC-region vibrations showed a decrease in amplitude and the tuning became almost low-pass but remained compressively nonlinear at all frequencies where responses were detectable. The response phases changed mildly. Amplitude recovery proceeded over several hours. 50 mins post-injection the BF-region BM and OHC vibrations remained depressed (Figs. 4 and 5I), while sub-BF OHC vibrations had fully recovered in expt. 785 (Fig. 4I), and substantially recovered in expt. 788 (Fig. 5I). 70 mins after the furosemide treatment, the BF-peak began to recover – the BF-peak recovery signifies the recovery of cochlear amplification (Figs. 4 and 5K). In expt. 785, both the BM and OHC vibrations had almost fully recovered at 120 mins (Fig. 4M), and in expt. 788, recovery was substantial at 120 mins (Fig. 5M). Beyond this time point the responses did not improve.

Prior to the furosemide injection, all the ears tested exhibited robust DPOAEs in response to swept two-tone stimuli. The 2 *f*_1_ – *f*_2_ levels were typically 30 - 40 dB lower than the primaries and present up to 32 kHz, the maximum frequency tested, when elicited with the 70 dB SPL primaries. With 50 dB SPL primaries they were measurably present to ~ 25 kHz. Just after furosemide, DPOAEs dropped into the noise level (~ 0 dB SPL) at all frequencies for the 50 dB primaries and at frequencies above ~ 15 kHz for the 70 dB primaries. Following an early burst of recovery at relatively low frequencies (below 10 kHz), DPOAEs gradually increased, showed their fullest recovery at ~ 120 mins, similar to the vibration. At that point DPOAEs had recovered fully at frequencies below ~ 22 kHz, which corresponds to *f*_2_ of ~ 32 kHz. The DPOAE recovery was not monotonic and the time course of the recovery supports the notion that operating point is a key component in the recovery of cochlear amplification, as will be discussed later.

Fig. 6 shows the time variation of the responses at BF and BF/2 for the two illustrative experiments. At BF/2, BM responses were unaffected by the furosemide treatment (Fig. 6C and G). In contrast, the OHC region BF/2 responses retained nonlinearity even immediately post-furosemide; the responses dropped slightly (at most a factor of ~4) and gain losses were largest at high SPL. The OHC-region BF/2 responses recovered nearly fully by 50 mins for expt. 785 (Fig. 6D); for expt. 788 the main recovery was by ~ 70 mins and was not full at high SPL (Fig. 6H). At BF, BM responses dropped most at low SPL and for both experiments at 40 and 50 dB SPL they dropped beneath the noise floor (Fig. 6A and E). There was considerable recovery at ~ 60–70 mins, when these low-SPL responses re-emerged, and continued recovery to 100 mins. The OHC-region BF responses showed an even greater disappearance into the noise (Fig. 6B and F), and by considering Figs. 4 and 5, this can be seen to be due to the low-pass characteristic of the post-furosemide OHC-region responses: responses cut off at frequencies below the BF. In the OHC-region, recovery of amplification at the BF was well underway at ~ 60 – 70 mins, as in the BM region, and continuing to 100 – 120 mins. The time course of amplification recovery as measured in these vibration responses is similar to what was observed in LCM BF responses (15). To illustrate that similarity, LCM BF data from Fig. 2B are plotted along with OHC-region curves in panels B and F. (The LCM data were scaled to overlie the vibration data, which does not affect the comparison since the y-axis is logarithmic.)

**Figure 6:**
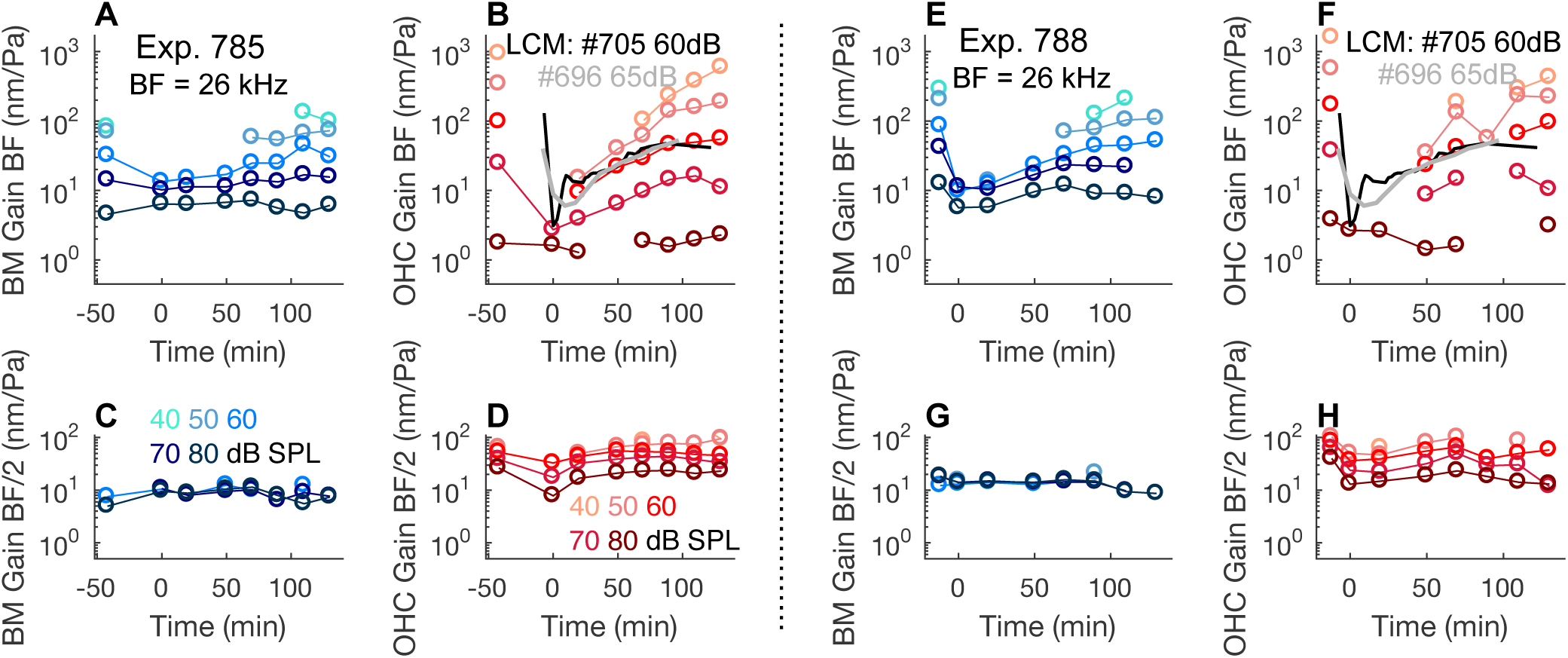
BM and OHC-region vibrations before, after, and during recovery from furosemide treatment, with time *t* = 0 right after the injection. Plots on the left are from the same experiment as Fig. 4; those on the right are from Fig. 5. (A) and (E) BM gains at the BF. (B) and (F) OHC-region gains at BF. (C) and (G) BM gains at BF/2. (D) and (H) OHC-region gains at BF/2. Gaps in the data occur when the responses are lower than the noise floor. The relatively long time between baseline and *t* = 0 in expt. 785 is present because, as noted in methods, in order to present the fullest (least noisy) baseline measurement data set, the set used for illustration was not always just before the injection. The grayscale data in (B) and (F) are BF LCM results from Fig. 2. These data are included to illustrate that BF LCM and vibration responses recovered on a similar time scale. The y-axis for the LCM data have units mV/Pa, and have been scaled by factors of 60 (expt. 705) and 40 (expt. 696).

In four preparations, recovery following furosemide never became substantial. Two examples are in Fig. 7. The reductions just post furosemide in these two preparations are similar to those in Figs. 4 and 5. In particular, the sub-BF nonlinearity is retained in these preparations just after furosemide, while amplification is absent. However, after faltering steps to recovery, amplification declined in these preparations. Fig. 8 contrasts the recovery of BF/2 responses averaged over the four animals with good recovery, and four animals with poor recovery. The initial drop in responses is similar, as is the initial recovery at 20 mins, but at 50 mins the "good" preparations are improving, while in the poor preparations recovery has turned into decline. The incomplete recovery in some preparations is likely due to a general decline in cochlear condition, which often happens in experimental cochlear physiology.

**Figure 7:**
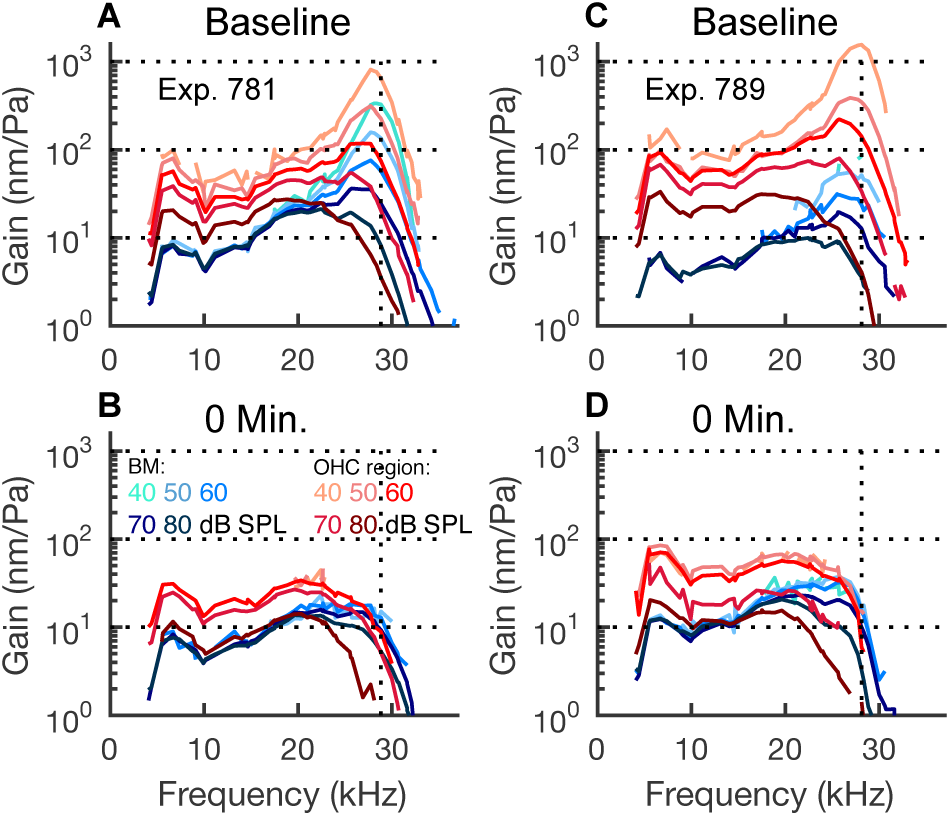
Loss of amplification following furosemide in two preparations that did not recover substantially. Gain relative to stimulus pressure at the ear canal. (A) Baseline, (B) Following furosemide, expt. 781. (C) Baseline, (D) Following furosemide, expt. 789.

**Figure 8:**
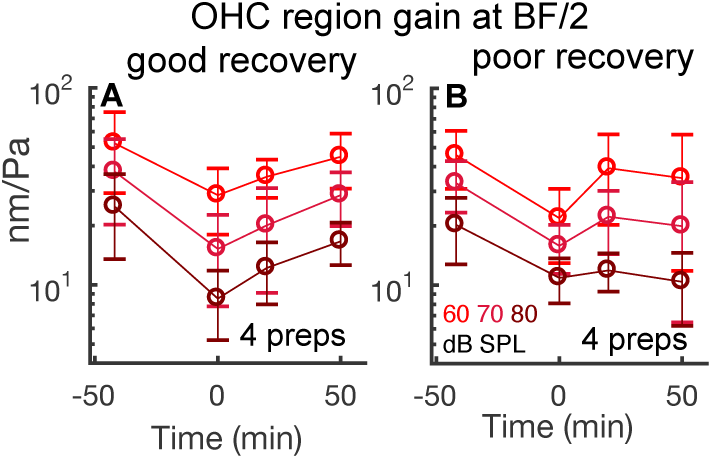
Reduction and recovery of OHC-region sub-BF responses following furosemide. Mean and standard deviation are shown for two groups of preparations. In (A) responses are tracked from the four preparations that showed substantial recovery of cochlear amplification (as in Figs. 4 and 5). In (B) responses are tracked from four preparations in which recovery faltered and never was substantial.

Returning to the preparations with good recovery, we consider the changes and recovery in the phase responses. Data are shown from the two experiments in Figs. 4 and 5 and a third experiment (783). The top row of Fig. 9 (A, C, E) shows baseline phase data and illustrates that BM and OHC-region phases go through a similar traveling wave excursion of several cycles. The second row (B, D, F) explores the phase differences, and how they change in time. In the baseline condition at low frequency (a few kHz) OHC-region vibration phase leads BM phase by up to ~ 0.2 cycle. This lead declines with frequency to eventually become a lag of ~ 0.1 cycle close to the BF. Just after furosemide, the OHC-BM phase difference dropped by ~ .05 cycle (18°) across frequency in the data of Fig. 9 B, and that observation approximately holds for the other two preparations in D and F. During the recovery period, that general-frequency drop in OHC–BM phase difference was partially but never fully reversed. In Fig. 9 G phase differences *within* each region before and just after furosemide are shown. This is grouped data, with mean and standard deviation from all eight preparations. The BM phase changed little and thus most of the time-dependent phase differences seen in Fig. 9 B, D and F resulted from changes in the OHC-region phase.

**Figure 9:**
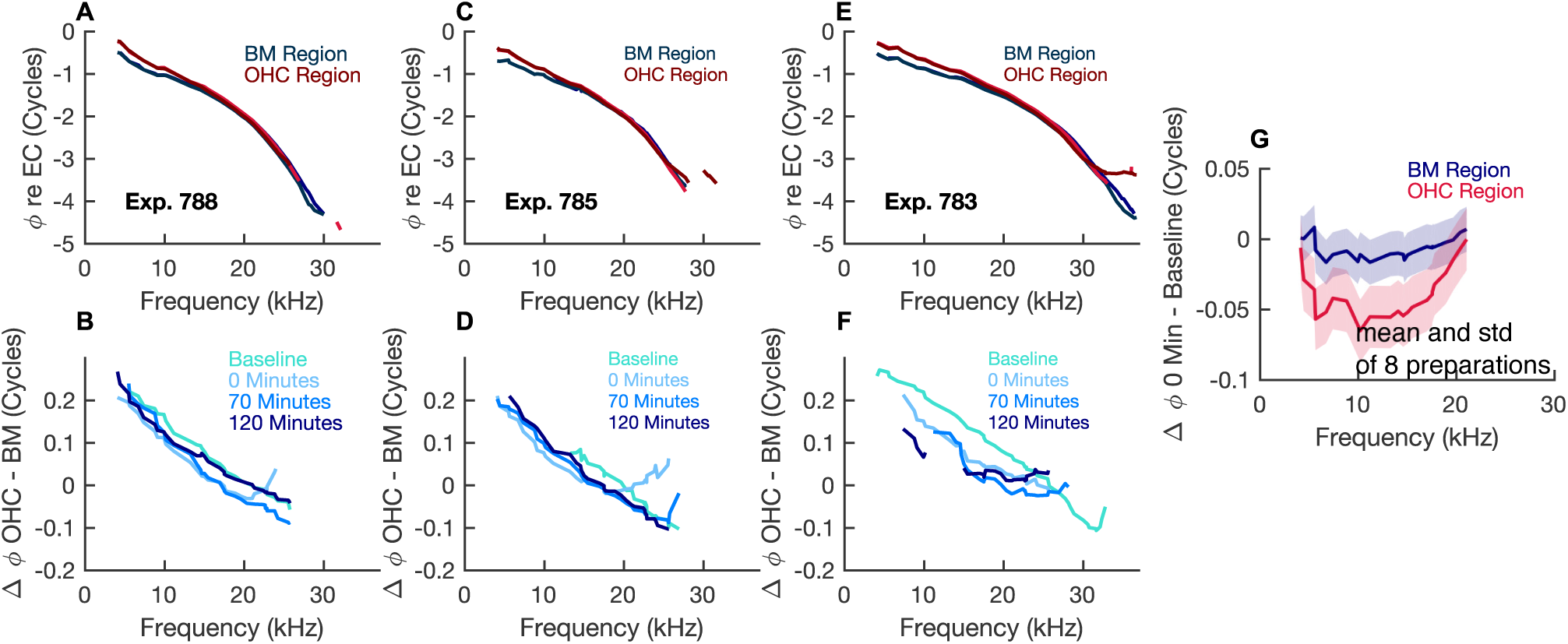
Expanded view of phase differences. Only relatively high SPL results are shown (70 and/or 80 dB SPL) because at lower SPL data often dropped beneath the noise floor following furosemide. (A, C, E) Subset of baseline phase data from expt.785 (Fig. 4), expt.788 (Fig. 5) and a third (expt.783) that showed substantial recovery. This row reinforces the basic similarity and small but robust differences between BM and OHC region phases at baseline. (B, D, F) Phase differences between OHC region and BM region, before and just after furosemide (t=0), at 120 min. when recovery was approximately fullest and at one intermediate time point (70 min). 70 dB SPL results shown. A three-point smoothing was done on these data to reduce distracting sharp variations. (G) Phase differences within each region before and just after furosemide. Mean and standard deviation from eight preparations are shown. This data is included to show that the BM-region phase changed little, and most of the time-dependent phase differences seen in (B, D, F) resulted from changes in OHC-region phase. 80 dB SPL results shown.

## DISCUSSION

The electrical gradient of the EP provides much of the driving force for the MET currents. In the OHCs, the receptor potentials generated by these currents drive electromotility which boosts BM and reticular lamina vibrations and is an essential component of the cochlea’s active process. We used furosemide to reduce EP and followed the recovery of OCC vibrations and DPOAEs over several hours.

BM and OHC-region vibrations recovered nearly fully in several preparations, and the time-scale of recovery in different frequency regions and in DPOAEs indicates that several processes are involved in recovery. OHC-region sub-BF vibration retained nonlinearity throughout, and was back to baseline by ~ 50 - 70 mins, signifying relatively rapid recovery of robust electromotility. The recovery of the BF peak in OHC-regions and at the BM, signifying functional cochlear amplification, occurred later, mainly during the time period between 70 and 120 mins. This two-stage recovery is reminiscent of the findings of Wang *et al.*, who studied changes in LCM following the same iv-furosemide protocol (15). In that study, EP had stabilized at a sub-normal level at ~ 40 mins and the LCM BF peak finally recovered at 100 mins or later. The timing of changes in LCM second harmonic responses indicated that the recovery of cochlear amplification occurred simultaneously with re-centering of the MET channel operating point. In the present study of OCC vibration, the comparable noise floor was higher than that of the LCM study, and second harmonics could not be examined. DPOAEs were used to explore operating point shifts in the present study.

The recovery of DPOAEs in Figs.4 and 5 is nonmonotonic, and a boost of recovery at ~ 70 mins is apparent. To observe this more fully, in Fig. 10 the recovery time course of an average of three DPOAEs at adjacent frequencies close to the BF of the measurement location are shown as line plots. Fig. 10 A and B are from the preparations in Figs. 4 and 5; Fig. 10 C is a mean of the four preparations in which amplification recovered most fully. The reduction at *t* = 0 corresponds to the dark line that appears at *t* = 0 in the heat plots of Figs. 4 and 5. Following that sharp reduction, the DPOAEs underwent 20 mins of recovery, but then dropped again, reached a local minimum at 50 mins and then entered a final recovery phase. The size of DPOAEs is expected to be directly affected by the reduction in EP, which would reduce OHC receptor current and voltage, and thus reduce active cochlear feedback. However, the nonmonotonic recovery illustrated in Fig. 10 cannot be attributed to the EP, which in our experience recovers monotonically and had likely stabilized at ~40 mins post-furosemide (15).

**Figure 10:**
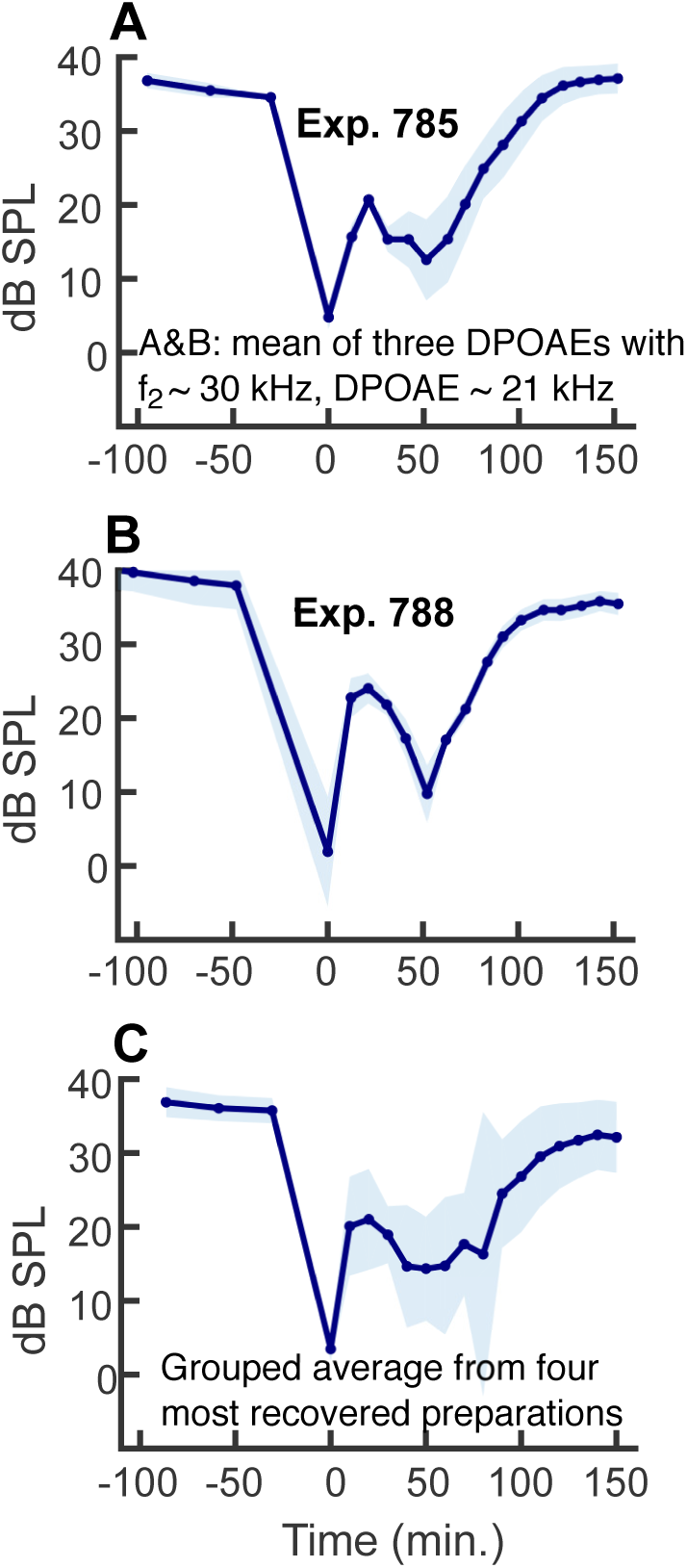
Recovery of DPOAEs. The three DPOAEs around *f*_2_ = 30 kHz, corresponding to DPOAE frequencies ~ 21 kHz were averaged and mean and standard deviation are shown. Primary levels were 70 dB SPL. (A) Expt. 785. (B) Expt. 788. (C) Results as in (A) and (B), averaged over the four preparations with substantial recovery.

These results can be considered through the lens of the MET operating point behavior found in (15). In Fig. 11 A, the MET function that was derived in (15) is shown with zero operating point offset, and with offsets of −0.12 and +.05 Pa. These offsets are included because in the findings shown in Fig. 2 C, the MET operating point was positive at a value of ~ .05 Pa at t=0, shifted negative to a minimum value of ~ −0.12 Pa at ~ 60 mins, and began to recenter (shift towards zero) at ~ 60 – 70 mins. Fig. 11 B shows how the size of the 2 *f*_1_ – *f*_2_ distortion produced by the nonlinearity in Fig. 11 A is affected by operating point shifts (7, 29). The primary inputs into the nonlinearity were at a level of 70 dB SPL (.06 Pa peak), to coincide with the primary inputs producing the DPOAEs in Fig. 10. Fig. 11 B shows that the 2 *f*_1_ − *f*_2_ distortion product will be a maximum with 0 offset, will reduce to zero at an offset of ~ ± .012 Pa, and increase for larger offsets. Thus, the second minimum in the DPOAE responses, starting its descent at ~ 25 mins in Fig. 10, and second boost of DPOAE recovery starting at ~ 50 mins, is likely due to MET operating point shift. The appearance of a local minimum in the DPOAE at 50 mins is predicted for an operating point that had shifted to its most negative value at 50 mins, and shifted back toward zero between 50 and 100 mins. It is confirming that this is the MET behavior that was observed in (15), redrawn in this paper’s Fig. 2 C. It is also worth underlining that between 50 and 100 mins is also when the BF peak – signifying cochlear amplification – underwent its steady recovery in BM and OHC-region responses.

**Figure 11:**
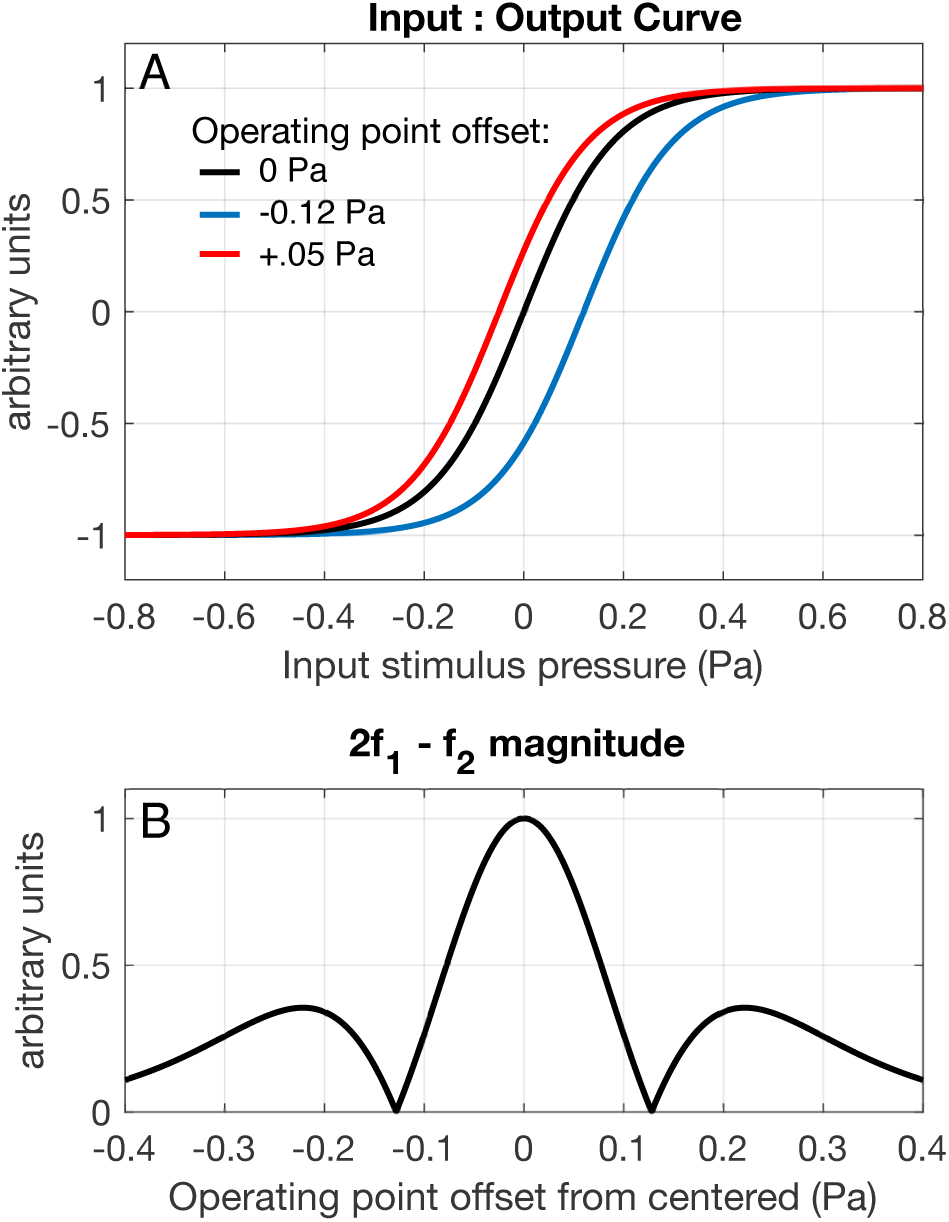
(A) A two-state Boltzmann function representing the nonlinear MET potential (*V*) as a function of the ear canal pressure (*P*) and operating point (*OP*), 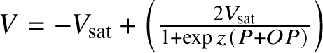. The slope factor *z* = −11.18 Pa^−1^, was derived in (15) based on LCM responses in the absence of amplification and the function has been plotted here on a normalized scale, with *V*_*sat*_ = 1. (The nonnormalized value for *V*_*sat*_ was 0.13 mV.) The OP shift that occurred after furosemide was shown in Fig. 2 C, and extended from −0.12 to +.05, and these extreme shifts are illustrated here. (B) Using the Boltzmann function, the size of the 2 *f*_1_ – *f*_2_ generated by this simple nonlinearity is found for a range of operating points. The input SPL of *f*_1_ and *f*_2_ was taken as 70 dB SPL (.06 Pa) to be consistent with Fig. 10.

This combination of findings focuses us on the questions: *What is required for cochlear amplification? What is essential, and what is sufficient?* We showed in (15) that a normal EP is not essential, and recovery of cochlear amplification at sub-normal EP had been hypothesized previously based on DPOAE recovery (16). OHC electromotility is essential based on genetic studies that changed prestin to a non-motile form (30). However, from the present study, the mere presence of robust electromotility is not sufficient for amplification of the BF-peak.

Simultaneous pressure and voltage measurements found that a phase shift between LCM and BM displacement was critical for OHC forces to be phased to provide amplification (19), and we explored the present data for time-dependence in the vibration phase following furosemide and recovery (Fig. 9 B, D, F.) As a broad-brush observation from these experiments, following furosemide the OHC-BM phase difference was offset by ~ -.05 cycle (-18°). This offset reduced the sub-BF phase lead of OHC-region relative to BM and increased the BF-region phase lag of OHC-region relative to BM. During the recovery of amplification, the most compelling phase recovery came from expt. 788 (Fig. 9 B) in which, in the 70-120 mins during which amplification recovered, this 18° furosemide-induced offset fully reversed at frequencies above 13 kHz. Thus, in this experiment the phasing between OHC and BM motion returned to a pre-furosemide character, concurrent with recovery of amplification. Expt. 785 also underwent a furosemide-induced negative offset in OHC-BM phase, and the offset recovered but only partially, although amplification showed substantial recovery. Expt. 783 also underwent a furosemide-induced negative offset in OHC-BM phase and in this case the offset recovered barely at all. In expt. 783, amplification recovery, while substantial, was less full than in the other two preparations. In sum, an OHC-BM phase change was robust following furosemide – a downward offset occurred between the baseline and *t* = 0 measurements in three of three explored data sets. Recovery of that offset was only weakly correlated with recovery of amplification. In the study of Cooper *et al.* (23) the phase of OHC vibration relative to BM was found to be viewing-angle dependent, and subtle preparation-viewing-angle differences could conceivably be affecting the repeatablity of phase changes in our study.

Based on the results in this study and its predecessor (15), the recovery of amplification occurs well after full recovery of electromotility and stabilization of EP, and concurrent with MET operating point recentering. These findings lend strength to the expectation that beyond the *size* of critical factors such as driving voltage and electromotility, amplification relies on qualitative conformational factors. That concept was illuminated by Jacob and Fridberger (31), who applied quasi-static current injections to restore the EP *in situ* in guinea pig temporal bone preparations. Using confocal fluorescent microscopy and interferometry they found that the organ of Corti underwent a series of mechanical conformational changes, shifting the apical region of the OHCs towards scala vestibuli, and these shifts positioned the system into a region of increased sensitivity. The position changes, ~ 50 – 100 nm, are smaller than the resolution of an OCT when used in two-dimensional imaging mode, so we cannot probe those findings directly with our imaging system. The recovery of amplification we observed occurred after EP was expected to be substantially recovered and stabilized, whereas the changes observed by Jacob *et al.* were concurrent with directly-induced EP changes. Despite this difference, the finding of that study, that conformational changes play a large role in setting the proper mechanical conditions for amplification, is supported by our findings.

Disrupting the EP could cause other perturbations that lead to operational changes that diminish auditory sensitivity, and these perturbations could recover on a different time scale than EP. In a recent review article, Manley (32) discusses the possibility that the EP exists at least in part to drive calcium metabolism in the mammalian cochlea, where evolutionary pressures caused the endolymphatic calcium concentration to fall to the micromolar level, a value significantly lower than in the auditory organs of other vertebrates. Calcium plays a number of roles in the hair cells including regulating the open probability of the MET channels, and may control one or more of the adaptation processes that partially set the dynamic range of the transduction process (33–35). Reduced EP will drive Ca^2+^ into scala media, which could affect the TM and its relationship to hair cells (36). In an *in situ* study, Strimbu *et al.* found that the calcium concentration in the TM is elevated relative to the bulk endolymph by ~ 25 *µ*M (37). In that study, following a brief exposure to loud sounds, both the TM calcium and cochlear microphonic were reduced and recovered in synchrony on a time-scale of ~ 30 mins. Using organ of Corti explants, Vélez-Ortega *et al.* showed that reducing the MET currents with channel blockers and manipulation of both intracellular and extracellular Ca^2+^ for as little as one hour could alter the morphology of the hair bundles. Reducing calcium entry into the bundles caused contraction of the shorter rows that contain the transduction channel and its associated machinery (38). If a similar effect occurs *in vivo* in an adult animal with fully mature hair cells, deleterious changes could occur to the hair bundles during the 30 – 40 mins following the furosemide injection when the EP remains significantly depressed. One would expect such physical changes to recover over a longer time scale than the recovery of EP.

In addition to reducing the EP, furosemide may reduce auditory sensitivity through other mechanisms. Loop diuretics weaken the blood labyrinth barrier by damaging the tight junctions lining the stria vascularis, an effect that has been exploited to enhance the uptake of contrast agents in MRI studies (39). The ototoxicity of other agents, such as cisplatin or aminoglycosides, is known to increase when administered simultaneously with loop diruetics, such as furosemide, an effect that presumably arises due to the increased permeability of the drugs in the perilymphatic spaces of the cochlea (40). Furosemide applied extracellularly to isolated outer hair cells diminishes nonlinear capacitance, a surrogate for electromotility (41). More generally, furosemide is known to affect the biophysical properties of lipids *in vitro* (42). Such changes to lipid bilayer of the hair cells’ bodies could alter electromotilty or the properties of the MET channels themselves (43, 44). However, the concentrations of furosemide in perilymph (45) and endolymph (46) following iv injections at the dosages used in this study have been measured and are in the micromolar range, approximately an order of magnitude lower than the extracellular concentration needed to evoke changes in nonlinear capacitance (41) or lipid properties *in vitro* (42). This suggests that any direct affects furosemide might have on the OHC bodies would be secondary in importance to the reduction of the EP.

In summary, we used the reversible changes caused by iv-furosemide to probe the essential ingredients of cochlear amplification. Following furosemide treatment the BF peak disappeared in BM and OHC-region mechanical responses while the sub-BF OHC-region nonlinearity was retained. Sub-BF OHC-region responses substantially or fully recovered to baseline levels during the first hour following furosemide. This suggests that OHC electromotility was fully operational at that point, and based on recent findings from our group, EP would have been stable at a sub-normal level. The BF peak underwent its own significant recovery during the second hour following furosemide, concurrent with an apparent re-centering of MET channels (15). These findings indicate that the presence of normal, high EP is not necessary for amplification and robust OHC electromotility is not by itself sufficient for cochlear amplification, and support the idea that the mechanical conformation of the organ of Corti complex is key to a functioning cochlear amplifier.

## AUTHOR CONTRIBUTIONS

C.E.S. Performed OCT experiments, analyzed the data and drafted the manuscript. Y.W. Performed iv injections in initial experiments, and contributed to study design. E.O. Supervised research, analyzed data and participated in manuscript writing.

### ACKNOWLEDGMENTS

This work was funded by NIH Grant R01-DC015362 and the Emil Capita Foundation.

## REFERENCES

1. Ashmore, J., 2008. Cochlear outer hair cell motility. Physiol. Rev. 88:173–210.

2. Robles, L., and M. A. Ruggero, 2001. Mechanics of the mammalian cochlea. Physiol. Rev. 81:1305–1352.

3. Hudspeth, A. J., 2008. Making an effort to listen: mechanical amplification in the ear. Neuron 59:530–545.

4. Ashmore, J., P. Avan, W. E. Brownell, P. Dallos, K. Dierkes, R. Fettiplace, K. Grosh, C. M. Hackney, A. J. Hudspeth, F. Julicher, B. Lindner, P. Martin, J. Meaud, C. Petit, J. Santos-Sacchi, J. R. Sacchi, and B. Canlon, 2010. The remarkable cochlear amplifier. Hear. Res. 266:1–17.

5. Peng, A. W., and A. J. Ricci, 2011. Somatic motility and hair bundle mechanics, are both necessary for cochlear amplification? Hear. Res. 273:109–122.

6. Fettiplace, R., and K. X. Kim, 2014. The physiology of mechanoelectrical transduction channels in hearing. Physiol. Rev. 94:951–986.

7. Brown, D. J., J. J. Hartsock, R. M. Gill, H. E. Fitzgerald, and A. N. Salt, 2009. Estimating the operating point of the cochlear transducer using low-frequency biased distortion products. J. Acoust. Soc. Am. 125:2129–2145.

8. Lukashin, A., and I. Russell, 1997. A descriptive model of the receptor potential nonlinearities generated by the hair cell mechanoelectric transducer. J. Acoust. Soc. Am. 103:973–980.

9. Keithley, E. M., 2019. Pathology and mechanisms of cochlear aging. J. Neurosci. Res. 0.

10. Tu, N. C., and R. A. Friedman, 2018. Age-related hearing loss: Unraveling the pieces. Laryngoscope Investig Otolaryngol 3:68–72.

11. Yamasoba, T., F. R. Lin, S. Someya, A. Kashio, T. Sakamoto, and K. Kondo, 2013. Current concepts in age-related hearing loss: epidemiology and mechanistic pathways. Hear. Res. 303:30–38.

12. Ruggero, M. A., and N. C. Rich, 1991. Furosemide alters organ of corti mechanics: evidence for feedback of outer hair cells upon the basilar membrane. J. Neurosci. 11:1057–1067.

13. Sewell, W. F., 1984. The effects of furosemide on the endocochlear potential and auditory-nerve fiber tuning curves in cats. Hear. Res. 14:305–314.

14. Komune, S., and T. Morimitsu, 1985. Dissociation of the cochlear microphonics and endocochlear potential after injection of ethacrynic acid. Arch Otorhinolaryngol 241:149–156.

15. Wang, Y., E. Fallah, and E. S. Olson, 2019. Adaptation of Cochlear Amplification to Low Endocochlear Potential. Biophys. J. 116:1769–1786.

16. Mills, D. M., S. J. Norton, and E. W. Rubel, 1993. Vulnerability and adaptation of distortion product otoacoustic emissions to endocochlear potential variation. J. Acoust. Soc. Am. 94:2108–2122. https://doi.org/10.1121/1.407483.

17. Schmiedt, R. A., H. Lang, H. O. Okamura, and B. A. Schulte, 2002. Effects of furosemide applied chronically to the round window: a model of metabolic presbyacusis. J. Neurosci. 22:9643–9650.

18. Fridberger, A., J. B. de Monvel, J. Zheng, N. Hu, Y. Zou, T. Ren, and A. Nuttall, 2004. Organ of Corti Potentials and the Motion of the Basilar Membrane. J of Neuroscience 24:10057–10063. https://www.jneurosci.org/content/24/45/10057.

19. Dong, W., and E. S. Olson, 2013. Detection of cochlear amplification and its activation. Biophys. J. 105:1067–1078.

20. Fallah, E., C. E. Strimbu, and E. S. Olson, 2019. Nonlinearity and amplification in cochlear responses to single and multi-tone stimuli. Hear. Res. 377:271–281.

21. Saremi, A., and S. Stenfelt, 2013. Effect of metabolic presbyacusis on cochlear responses: a simulation approach using a physiologically-based model. J. Acoust. Soc. Am. 134:2833–2851.

22. Lin, N. C., C. E. Strimbu, C. P. Hendon, and E. S. Olson, 2018. Adapting a commercial spectral domain optical coherence tomography system for time-locked displacement and physiological measurements. AIP Conference Proceedings 1965:080004.

23. Cooper, N. P., A. Vavakou, and M. van der Heijden, 2018. Vibration hotspots reveal longitudinal funneling of sound-evoked motion in the mammalian cochlea. Nat Commun 9:3054.

24. Versteegh, C. P., and M. van der Heijden, 2012. Basilar membrane responses to tones and tone complexes: nonlinear effects of stimulus intensity. J. Assoc. Res. Otolaryngol. 13:785–798.

25. Chen, F., D. Zha, A. Fridberger, J. Zheng, N. Choudhury, S. L. Jacques, R. K. Wang, X. Shi, and A. L. Nuttall, 2011. A differentially amplified motion in the ear for near-threshold sound detection. Nat. Neurosci. 14:770–774.

26. Lee, H. Y., P. D. Raphael, A. Xia, J. Kim, N. Grillet, B. E. Applegate, A. K. Ellerbee Bowden, and J. S. Oghalai, 2016. Two-Dimensional Cochlear Micromechanics Measured In Vivo Demonstrate Radial Tuning within the Mouse Organ of Corti. J. Neurosci. 36:8160–8173.

27. Ren, T., W. He, and D. Kemp, 2016. Reticular lamina and basilar membrane vibrations in living mouse cochleae. Proc. Natl. Acad. Sci. U.S.A. 113:9910–9915.

28. Lin, N. C., C. P. Hendon, and E. S. Olson, 2017. Signal competition in optical coherence tomography and its relevance for cochlear vibrometry. J. Acoust. Soc. Am. 141:395–405. https://doi.org/10.1121/1.4973867.

29. Sirjani, D. B., A. N. Salt, R. M. Gill, and S. A. Hale, 2004. The influence of transducer operating point on distortion generation in the cochlea. J. Acoust. Soc. Am. 115:1219–1229.

30. Dallos, P., X. Wu, M. A. Cheatham, J. Gao, J. Zheng, C. T. Anderson, S. Jia, X. Wang, W. H. Cheng, S. Sengupta, D. Z. He, and J. Zuo, 2008. Prestin-based outer hair cell motility is necessary for mammalian cochlear amplification. Neuron 58:333–339.

31. Jacob, S., M. Pienkowski, and A. Fridberger, 2011. The endocochlear potential alters cochlear micromechanics. Biophys. J. 100:2586–2594.

32. Manley, G. A., 2017. The mammalian Cretaceous cochlear revolution. Hear. Res. 352:23–29. http://www.sciencedirect.com/science/article/pii/S0378595516305275, annual Reviews 2017.

33. Peng, A. W., T. Effertz, and A. J. Ricci, 2013. Adaptation of mammalian auditory hair cell mechanotransduction is independent of calcium entry. Neuron 80:960–972.

34. Corns, L. F., S. L. Johnson, C. J. Kros, and W. Marcotti, 2014. Calcium entry into stereocilia drives adaptation of the mechanoelectrical transducer current of mammalian cochlear hair cells. Proc. Natl. Acad. Sci. U.S.A. 111:14918–14923. https://www.pnas.org/content/111/41/14918.

35. Caprara, G. A., A. A. Mecca, Y. Wang, A. J. Ricci, and A. W. Peng, 2019. Hair Bundle Stimulation Mode Modifies Manifestations of Mechanotransduction Adaptation. J. of Neurosci. 39:9098–9106.

36. Freeman, D. M., K. Masaki, A. R. McAllister, J. L. Wei, and T. F. Weiss, 2003. Static material properties of the tectorial membrane: a summary. Hear. Res. 180:11–27. http://www.sciencedirect.com/science/article/pii/S0378595503000728.

37. Strimbu, C. E., S. Prasad, P. Hakizimana, and A. Fridberger, 2019. Control of hearing sensitivity by tectorial membrane calcium. Proc. Natl. Acad. Sci. U.S.A. 116:5756–5764. https://www.pnas.org/content/116/12/5756.

38. Vélez-Ortega, A. C., M. J. Freeman, A. A. Indzhykulian, J. M. Grossheim, and G. I. Frolenkov, 2017. Mechanotransduction current is essential for stability of the transducing stereocilia in mammalian auditory hair cells. eLife 6:e24661. https://doi.org/10.7554/eLife.24661.

39. Videhult Pierre, P., J. E. Rasmussen, S. Nikkhou Aski, P. Damberg, and G. Laurell, 2020. High-Dose Furosemide Enhances the Magnetic Resonance Signal of Systemic Gadolinium in the Mammalian Cochlea. Otol. Neurotol. 41:545–553. https://www.overleaf.com/project/566b68aa1d7510001698d90.

40. Ding, D., H. Liu, W. Qi, H. Jiang, Y. Li, X. Wu, H. Sun, K. Gross, and R. Salvi, 2016. Ototoxic effects and mechanisms of loop diuretics. J. Otol. 11:145–156.

41. Santos-Sacchi, J., M. Wu, and S. Kakehata, 2001. Furosemide alters nonlinear capacitance in isolated outer hair cells. Hear. Res. 159:69 – 73. http://www.sciencedirect.com/science/article/pii/S0378595501003215.

42. Bach, D., C. Vinkler, I. R. Miller, and S. R. Caplan, 1988. Interaction of furosemide with lipid membranes. J. Membr. Biol. 101:103–111.

43. Peng, A. W., R. Gnanasambandam, F. Sachs, and A. J. Ricci, 2016. Adaptation Independent Modulation of Auditory Hair Cell Mechanotransduction Channel Open Probability Implicates a Role for the Lipid Bilayer. J. Neurosci. 36:2945–2956.

44. Gianoli, F., T. Risler, and A. S. Kozlov, 2017. Lipid bilayer mediates ion-channel cooperativity in a model of hair-cell mechanotransduction. Proc. Natl. Acad. Sci. U.S.A. 114:E11010–E11019. https://www.pnas.org/content/114/51/E11010.

45. Rybak, L. P., T. P. Green, S. K. Juhn, T. Morizono, and B. L. Mirkin, 1979. Elimination Kinetics of Furosemide in Perilymph and Serum of the Chinchilla: Neuropharmacologic Correlates. Acta Oto-Laryngologica 88:382–387.

46. Hara, A., K. Machiki, M. Senarita, M. Komeno, and J. Kusakari, 1993. Pharmacokinetics of furosemide in endolymph. Auris Nasus Larynx 20:247–254.

